# CD40 Expression by B cells is Required for Optimal Immunity to Murine *Pneumocystis* Infection

**DOI:** 10.1101/2024.02.05.578900

**Authors:** Monica Sassi, Shelly J. Curran, Lisa R. Bishop, Yueqin Liu, Joseph A. Kovacs

## Abstract

CD40-CD40L interactions are critical for controlling *Pneumocystis* infection. However, which CD40-expressing cell populations are important for this interaction have not been well-defined. We used a cohousing mouse model of *Pneumocystis* infection, combined with flow cytometry and qPCR, to examine the ability of different populations of cells from C57BL/6 mice to reconstitute immunity in CD40 knockout (KO) mice. Unfractionated splenocytes, as well as purified B cells, were able to control *Pneumocystis* infection, while B cell depleted splenocytes and unstimulated bone-marrow derived dendritic cells (BMDCs) were unable to control infection in CD40 KO mice. *Pneumocystis* antigen-pulsed BMDCs showed early, but limited, control of infection. Consistent with recent studies that have suggested a role for antigen presentation by B cells, using cells from immunized animals, B cells were able to present *Pneumocystis* antigens to induce proliferation of T cells. Thus, CD40 expression by B cells appears necessary for robust immunity to *Pneumocystis*.

## Introduction

*Pneumocystis jirovecii* causes life-threatening disease, *Pneumocystis* pneumonia (PCP), in immunocompromised individuals [1, 2]. Although historically associated with HIV/AIDS patients, PCP also affects individuals receiving immunosuppressive therapy for a variety of medical conditions, as well as individuals with congenital immunodeficiencies, especially severe combined immunodeficiency (SCID) or X-linked hyper-IgM syndrome, which is due to a defect in CD40-CD40L interactions [3–6].

There are multiple *Pneumocystis* species, each of which appears to infect one or a limited number of closely related mammalian species [7]. Mouse models of *P. murina* infection have provided important insights into relevant host-organism interactions [8]. *Pneumocystis* enters the host via the respiratory tract and attaches primarily to type I alveolar epithelial cells [9]. Antigen-presenting cells presumably take up and process *Pneumocystis* antigens, such as surface glycoproteins, then migrate to peripheral lymphoid organs and present antigen to CD4^+^ T cells via the MHC II complex. For a robust T cell response, multiple interactions with costimulatory molecules need to occur between a T cell and an antigen presenting cell, resulting in T cell activation.

CD4^+^ T cells are essential for clearing *Pneumocystis*, and CD40-CD40L interactions appear to play a critical role [10–14]. We and others have previously demonstrated that infection of CD40 knockout (CD40 KO) or CD40L knockout (CD40L KO) mice with *P. murina* results in uncontrolled infection and eventual death, supporting the importance of these interactions in clearing *Pneumocystis* infection [12, 14, 15]. Microarray studies from our group have demonstrated largely absent immune responses in CD40L KO mice at a time that robust responses developed in immunocompetent mice following infection, suggesting an early role for the CD40-CD40L interaction in controlling infection [13].

CD40L is expressed predominantly on activated T cells, but during inflammation can also be expressed by activated B cells, monocytes, basophils, mast cells, natural killer cells, and platelets [16, 17]. CD40 is expressed primarily by B cells, dendritic cells and macrophages, all of which can function as antigen presenting cells (APCs) [16, 17]. B cells have been increasingly found to play a critical role in host immune responses to *Pneumocystis*, potentially for their role as APCs in addition to their role as antibody producers [18–22].

Given the importance of CD4^+^ T cells and the CD40-CD40L interaction to the control and clearance of *Pneumocystis*, we sought to determine, in a mouse model, which cell types expressing CD40 are the most critical to generating an effective immune response against *Pneumocystis* infection.

## METHODS

### Animals

Healthy C57BL/6 mice (expressing CD45.2) were obtained from the National Cancer Institute. CD40 ligand knock-out mice, (CD40L KO, strain B6, 129S-Tnfsf5^tm1lmx^/J), CD40 knock-out mice (CD40 KO, strain B6, 129P2-Cd40^tm1Kik/^J) and C57BL/6 mice expressing CD45.1 (B6.SJL-Ptprc^a^ Pepc^b^/BoyJ) were obtained from Jackson Laboratory (Bar Harbor, ME). All mouse strains were subsequently bred at the NIH Clinical Center animal facility, Bethesda, Maryland. Mice were housed in microisolator cages and kept in ventilated racks. All animal work was performed under an animal study protocol approved by the NIH Clinical Center Office of Animal Care and Use.

### Experimental design

We utilized a model of *Pneumocystis* infection that simulates naturally acquired infection [13, 15, 23]. CD40 KO mice were co-housed with a *P. murina*-infected CD40L KO seeder mouse, with a maximum of 11 animals per cage. Cells from naïve C57BL/6 mice, expressing CD45.1 as a congenic marker, were injected by tail-vein, as described below, into recipient CD40 KO mice expressing CD45.2 at ∼20-27 days after the start of cohousing. At ∼35, 65 and 90 days post-initial exposure, CD40 KO mice were sacrificed and spleens, blood, and lungs were collected. Because of the limited numbers of mice that could be housed per cage, and the need to co-house mice that received cells together with mice that received only saline, to ensure similar exposure to the seeders, data from multiple replicate cages were combined.

### Adoptive transfer of splenic cells and B cells

Immune reconstitution of CD40 KO mice was performed by tail vein injection. Briefly, spleens were collected from naïve C57BL/6 (CD45.1^+^) and a single cell suspension was obtained by incubation with Liberase Blendzyme 3 (Roche, Bradford, CT) and DNase (Sigma, St. Louis, MO) for 10 minutes at 37°C, followed by washing and removal of erythrocytes by ACK lysis buffer. Cells were counted and re-suspended in PBS, and splenocytes (∼50 x10^6^/mouse in 300 μl) were injected by tail vein into CD40 KO mice that had been exposed to *P. murina* for ∼3-4 wks.

CD19^+^ B cells were purified by positive magnetic separation using CD19 MicroBeads (Miltenyi Biotec, Gaithersburg, MD) and MS or LS columns, according to the manufacturer’s instructions. For CD19-depleted cells, the unlabeled cells that passed through the column during CD19^+^ cell purification were collected and further purified by application onto an LD column (Miltenyi Biotech). Cells derived from 3-4 mice were utilized in order to inject ∼30-50x10^6^ CD19^+^ cells/mouse or 15-25 x10^6^ CD19-cells/mouse in 300 µl PBS. Control mice were injected with 300μl of PBS alone.

Because tail vein injection was not always successful, and CD45.1^+^ cells were easily detectable in most mice that received splenic cells, data from mice scheduled to receive cells that had no detectable CD45.1^+^ cells in the lungs or spleen were excluded from analysis.

### Adoptive transfer of dendritic cells

Femurs collected from C57BL/6 WT mice expressing CD45.1 were flushed to obtain bone marrow cells, which were washed twice, then incubated in complete differentiation medium (RPMI-1640 supplemented with 10% heat-inactivated fetal bovine serum (FBS), 100U/ml penicillin, 100 μg/ml streptomycin, 5 mM glutamine (Invitrogen, Waltham, MA), 50 µM 2-ME (Sigma) and 30 ng/ml GM-CSF (Peprotech, Cranbury, NJ)), which was replaced every 2-3 days, to generate bone marrow-derived dendritic cells (BMDCs) [24, 25]. Non-adherent and loosely adherent cells were collected after 7-8 days and the immature BMDCs were magnetically sorted based on the expression of CD11c. Briefly, the cells were counted and incubated with mouse CD11c magnetic beads (Miltenyi, Biotec) for 15 minutes at 4°C. After washing, the cells were applied to LS columns with a pre-filter (Miltenyi, Biotech) and magnetically sorted following the manufacturer’s instructions. Eluted CD11c^+^ cells were washed and then either rested or primed with 20 µg/ml crude *P. murina* antigen in complete differentiation medium overnight at 37°C. For antigen preparation, *Pneumocystis* organisms were partially purified by Ficoll-Hypaque density gradient centrifugation [26], then disrupted with glass beads followed by sonication; the supernatant remaining after centrifugation was utilized as the antigen. BMDCs were collected and re-suspended in 300 µl of PBS. Each CD40 KO mouse received ∼ 2.5-10 x 10^6^ BMDCs, or PBS alone by tail vein injection. An aliquot was saved for flow cytometry analysis to verify the maturation stage of the BMDCs. Because no CD45.1^+^ cells were detected in mice receiving BMDCs, all mice were included in the analyses.

### Quantification of P. murina organism load

DNA was extracted from mouse lung tissue (stored at -80°C), using the QIAamp DNA Mini Kit (Qiagen, Germantown, MD) according to the manufacturer’s protocol. Intensity of *P. murina* infection was quantitated by qPCR using a single-copy gene (*dhfr*), as previously described [15]; organism loads are expressed as *dhfr* copies per mg lung tissue.

### Detection of anti-P. murina antibodies by ELISA

Serum anti-*P. murina* antibodies were detected by ELISA using a crude *P. murina* antigen, prepared as noted above, as previously described [27]. An optical density > 0.2 after subtracting background was used as a cutoff for a positive antibody response.

### Flow cytometry analysis of lung and spleen

Freshly removed lungs were incubated with collagenase (Gibco, Waltham, MA) and DNase (Sigma) at 37°C for 30 minutes to obtain single cell suspensions. After washing the cells with RPMI-10% FCS (Invitrogen), erythrocytes were removed with ACK lysis buffer and cells were washed twice more. Spleens were processed as described above to obtain single cells suspension. The cell suspensions (lung or spleen) were filtered with cell strainer, counted, and stained with LIVE/DEAD fixable Near-IR or violet dead cell kit according to the manufacturer’s protocol (Invitrogen). The cells were stained with the following fluorochrome-conjugated anti-murine antibodies: CD3-FITC or CD3-PE, CD4-PerCP, CD8-PerCP, CD19-PE or CD19-APC, CD11c-PE, CD11b-PerCP, CD45.1-PerCP-Cy5.5, and CD45.2 APC (all BD Biosciences, Franklin Lakes, NJ) and run on a FACSCanto (BD Biosciences); data were analyzed using FlowJo (BD Life Sciences, Ashland, OR).

### *In vitro* antigen presentation assays

To evaluate antigen presentation in vitro, mice were immunized with 20 µg of a crude *P. murina* antigen, prepared as noted above, together with Freund’s complete (first immunization) or incomplete (booster immunizations) adjuvant; 3 booster immunizations were administered approximately 2 weeks apart. Mice immunized with adjuvant alone served as controls. Spleen cells were processed as above, and cells from 3-4 mice were combined for sorting by flow cytometry, which was performed by the NHLBI core flow cytometry facility. The following fluorochrome-conjugated anti-murine antibodies: CD3-FITC, CD4-PE-Cy7, CD19-PE, CD11c-APC, and CD11b-APC (all BD Biosciences) together with LIVE-DEAD violet dead cell stain (Invitrogen) were utilized for sorting. Sorted CD4^+^ T cell and B cell purity was >95%, and DC/monocyte (CD11b+ and/orCD11c^+^) purity was ∼85%, as confirmed by flow cytometry. To evaluate antigen presentation to CD4^+^ T cells, CD4^+^ T cells (CD3^+^CD4^+^) were cultured either alone or with B cells (CD19^+^) or combined DCs/monocytes (CD11c^+^ and/or CD11b^+^) in the presence of 20 µg/ml of a crude *P. murina* antigen, 10 µg/ml of purified *P. murina* major surface glycoprotein (Msg) prepared as previously reported [27], or no antigen, for 5 days, after which proliferation was measured using CellTiter-Glo Luminescent Cell Viability Assay (Promega, Madison, WI) according to the manufacturer’s instructions. Proliferation results are presented as the stimulation index (mean proliferation, in relative light units (RLU), for antigen co-incubation divided by the mean RLU for the no antigen control).

### Statistics

For cell transfer studies, organism loads at individual time-points for mice receiving cell transfers were compared to organism loads in control mice, and for in vitro proliferation studies, the RLU for cells with antigen co-incubation were compared to the RLU for the no antigen control, using unpaired, 2-sided Student’s T-test, using Prism (GraphPad Software, LLC, San Diego, CA) or Excel (Microsoft, Redmond, WA).

## RESULTS

### Spleen cells from wild type mice can reduce *Pneumocystis* burden in the lungs of CD40 KO mice

To determine whether reconstituting the CD40-CD40L interaction in immune cells is enough to control *Pneumocystis* infection, we adoptively transferred splenic cells from naïve wild-type C57BL/6 mice into CD40 KO mice after approximately 3 weeks of continuous co-housing with a *P. murina*-infected seeder. In preliminary experiments, transferring splenic cells simultaneously with initial exposure to the seeder did not impact infection, and no CD45.1^+^ cells were detected in the lungs of reconstituted mice. CD45.1^+^ cells were easily detected in both the lungs and spleen of mice following cell transfer at 3 weeks (Figure 1).

**Figure 1.**
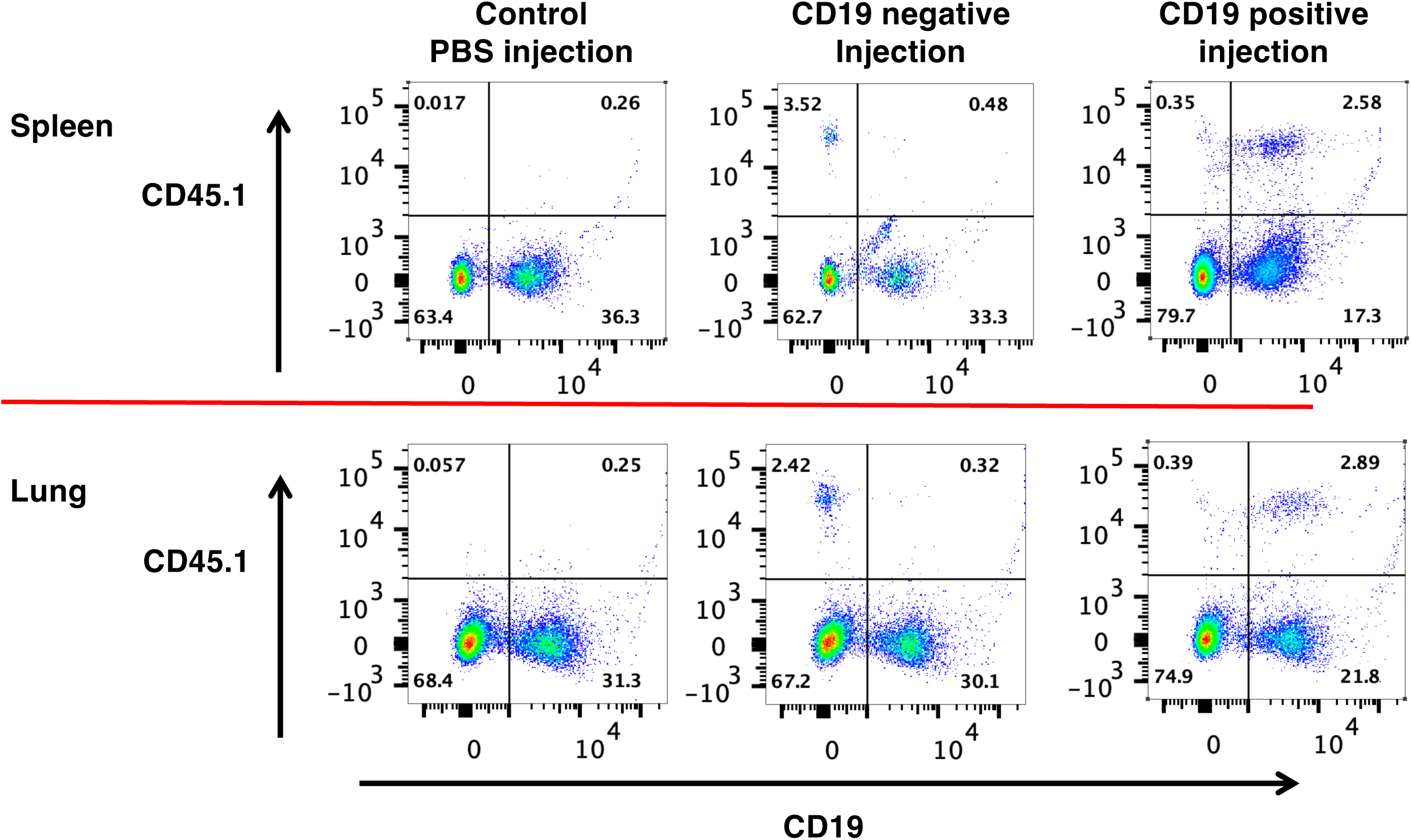
Flow cytometry analysis of lung and spleen cells following transfer of CD19-cells or CD19^+^ B cells; gating was on live cells, then lymphocytes, then CD19 and CD45.1. Representative results are shown for spleen and lung cells from CD40 KO mice exposed to a *Pneumocystis*-infected seeder for 65 days. The exposed mice were reconstituted at day 20 with PBS (control) or with CD19^+^ B cells (positive), or with CD19 depleted (negative) spleen cells. Transferred B cells (CD45.1^+^/CD19^+^) were detected in the spleen and lung of the mouse that received CD19+ B cells, while transferred non-B cells were detected in the mouse that received cells that had been depleted of CD19^+^ B cells. Neither population was detected in the mouse that had received PBS.

By 65 days post initial exposure, CD40 KO mice receiving wild-type splenic cells had a trend toward reduced *P. murina* compared to mice receiving saline (Figure 2). By 90 days, *P. murina* burden was significantly reduced (p=0.02) in the group receiving splenic cells compared to controls. As noted above, transferred CD45.1^+^ cells were detected in the spleen as well as in the lungs of recipient animals (Figure 1), and anti-*Pneumocystis* antibodies were detected in splenic cell recipients but not controls by days 65 and 90 (data not shown). These data demonstrate that reconstitution of the CD40-CD40L interaction by splenic cells protects CD40 KO mice from progressive *P. murina* infection.

**Figure 2.**
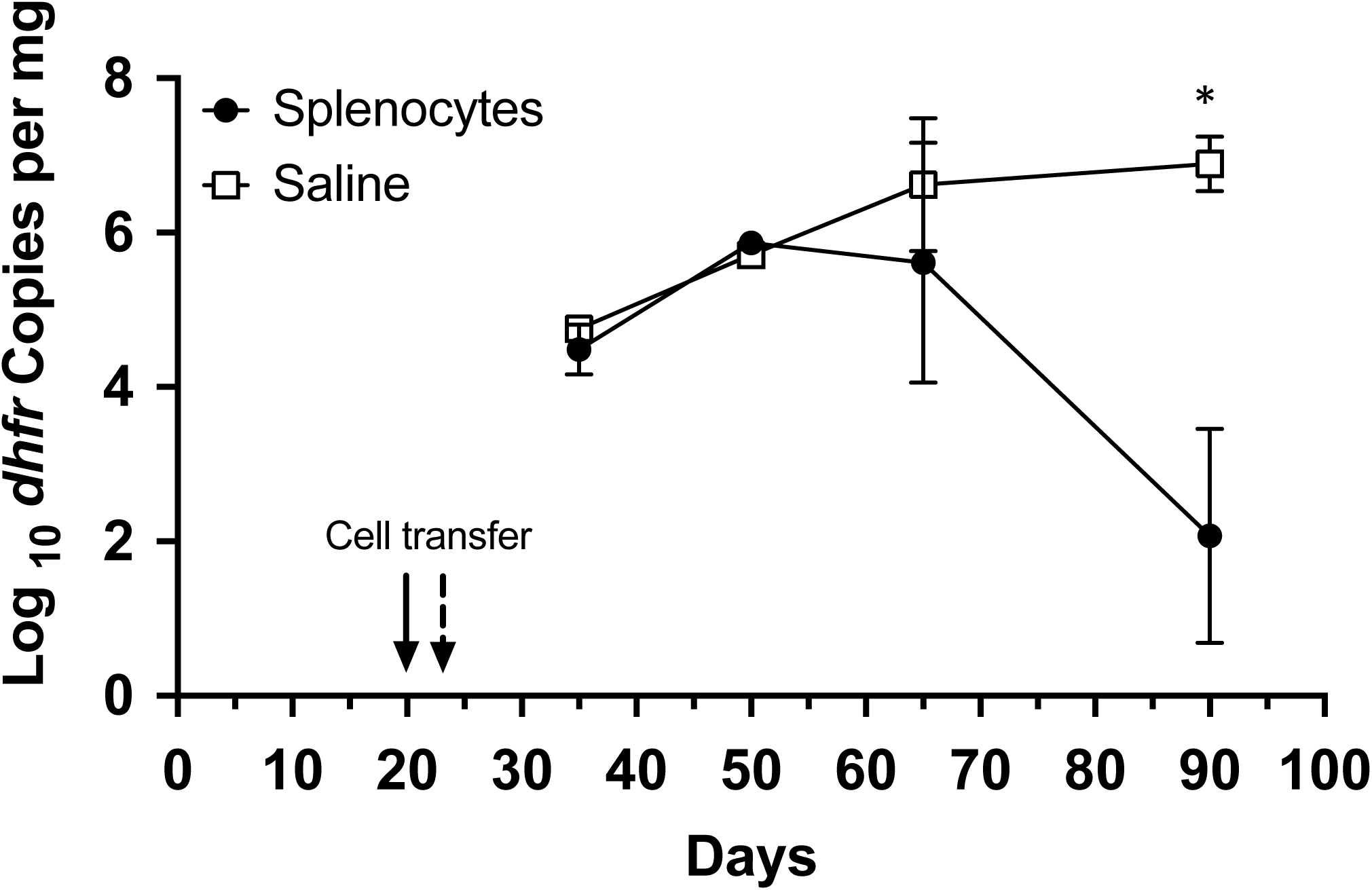
Transfer of splenocytes controls *P. murina* infection in CD40 KO mice. CD40 KO mice were cohoused with a *P. murina* infected seeder beginning at day 0. At day 20 (Cage 1) or 24 (cage 2) mice were injected via tail-vein with either PBS or ∼50 million splenocytes from C57BL/6 mice. Organism loads were deteremined by qPCR targeting the single copy *dfhr* gene, and are expressed as *dhfr* copies per mg lung tissue. Data represent the geometric mean ± SD for 2-4 mice per time-point for the splenocyte group, and 1 (day 50) to 2 mice per time-point for the PBS control group. P value is shown for the difference between the 2 groups, using Student’s unpaired t-test. *, p<0.05

### CD19^+^ B cell transfer is sufficient to reduce *Pneumocystis* burden in CD40 KO mice

Recently, B cells were shown to have potential additional roles, aside from antibody production, in controlling *Pneumocystis* infection, including antigen presentation, resulting in priming of CD4^+^ T cells and regulating the T helper response [19–22, 28]. Transfer of wild-type CD19^+^ B cells alone was able to control infection, with a nearly 4 log decline in organism load by day 90 compared to saline controls (Figure 3A). To explore this further, we adoptively transferred wild-type CD19^+^ cells or CD19^-^ cells into CD40 KO mice approximately three weeks after initial exposure to *P. murina*; by 65 days post initial exposure, CD19^+^ B cells again resulted in a significant decline in *Pneumocystis* burden compared to either CD19^-^ cells or saline (Figure 3B). This difference was more pronounced by 90 days post infection, with a nearly 3 log decline. Donor cells (CD45.1^+^) could be detected in both the lungs and spleens of mice that received either CD19^+^ or CD19^-^ cells but not in the mice that received saline (Figure 1). No differences in CD4^+^ or CD8^+^T cell numbers were seen among the different groups at either day 65 or day 90 (Figure 3C). Furthermore, an increase in serum anti-*Pneumocystis* Ig levels was present in the CD40 KO mice that received CD19^+^ cells but not in those that received CD19^-^ cells or saline alone, confirming reconstitution of an anti-*P. murina* humoral response (Figure 3D). Together, these data demonstrate that CD19^+^ B cells play a major role in controlling and clearing *Pneumocystis* infection via a mechanism mediated by the CD40-CD40L interaction, while CD19^-^ cells, which would include DCs, macrophages, CD4^+^ and CD8^+^ T cells, as well as other cell types, were unable to control infection.

**Figure 3.**
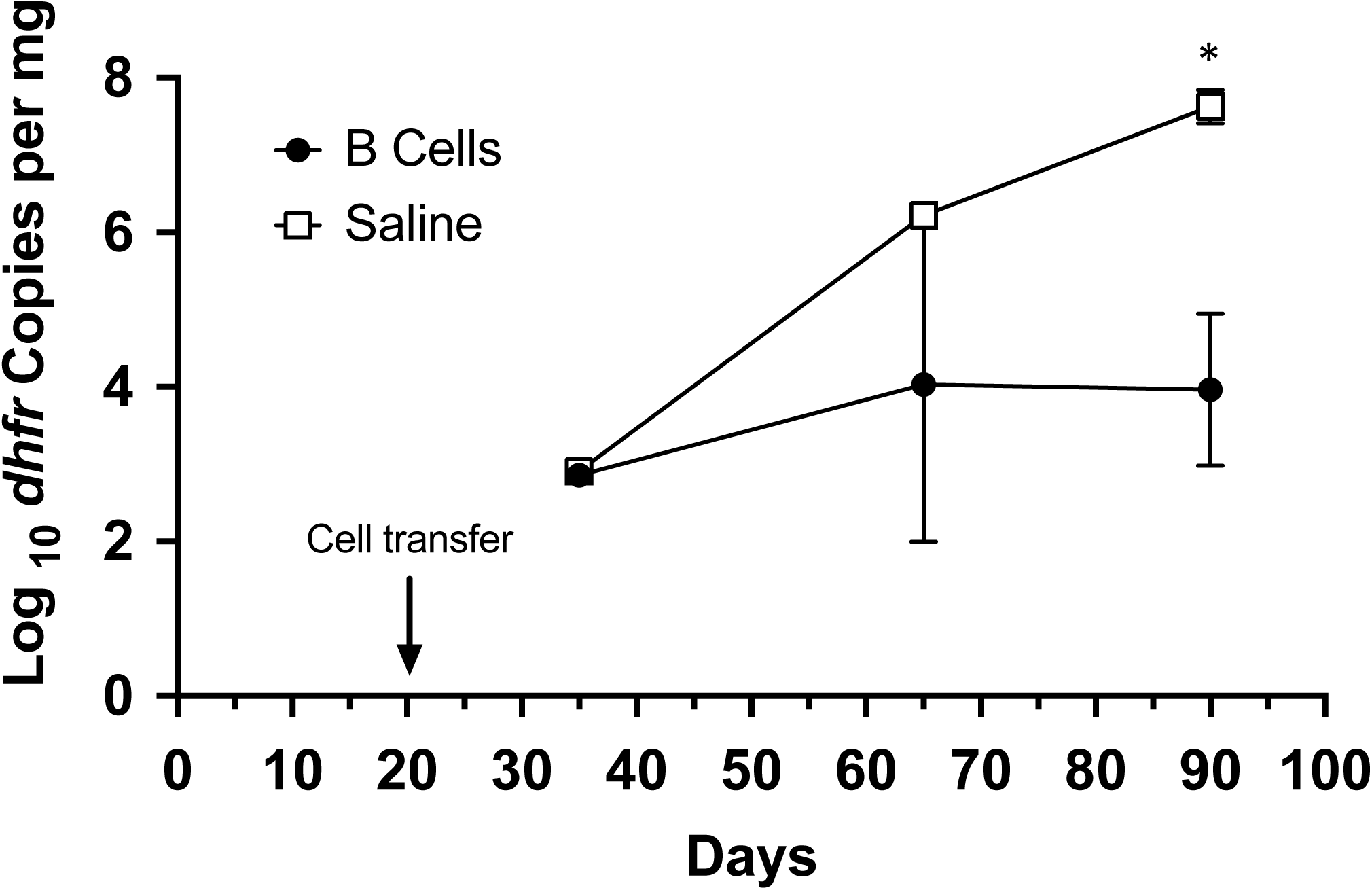
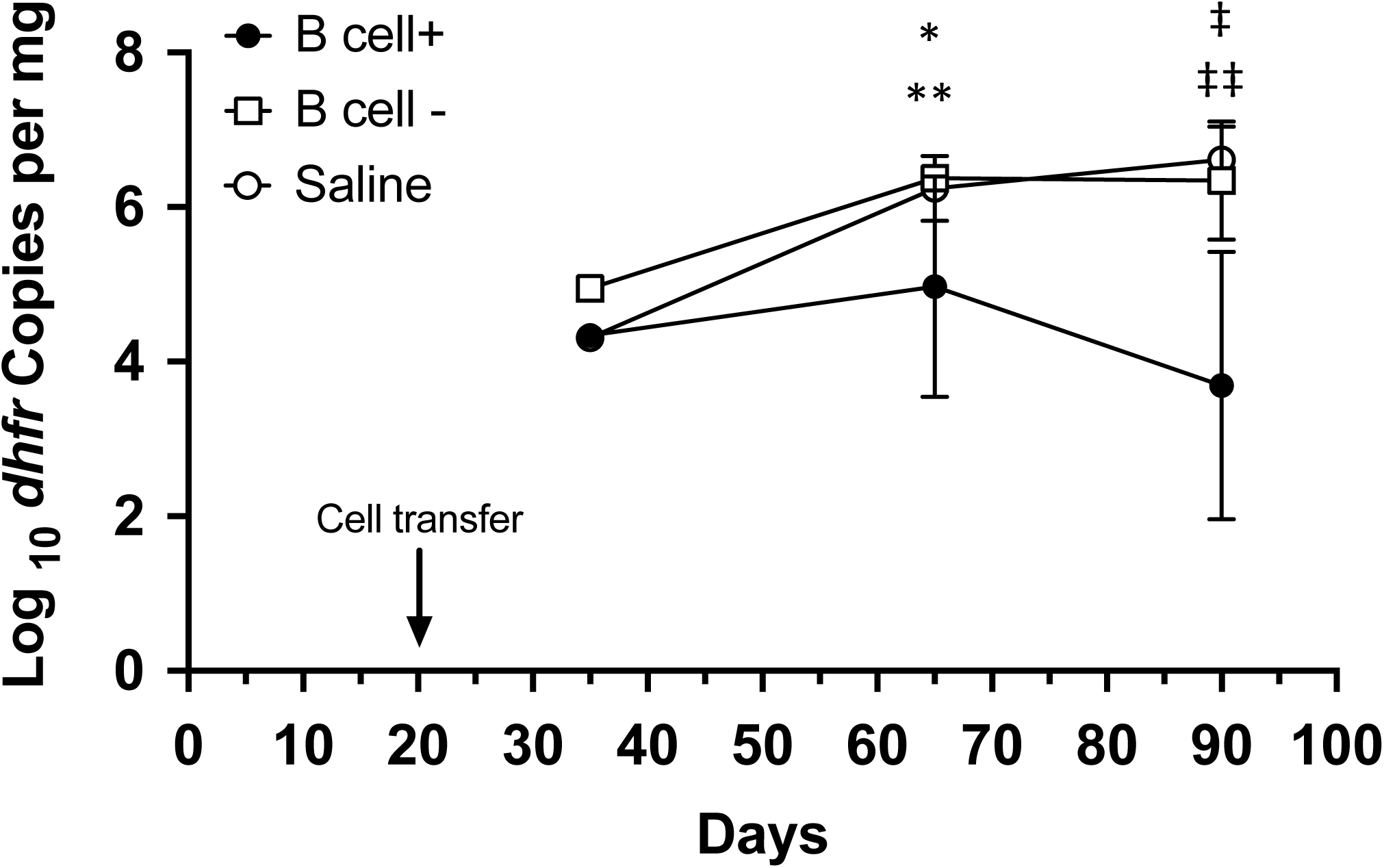
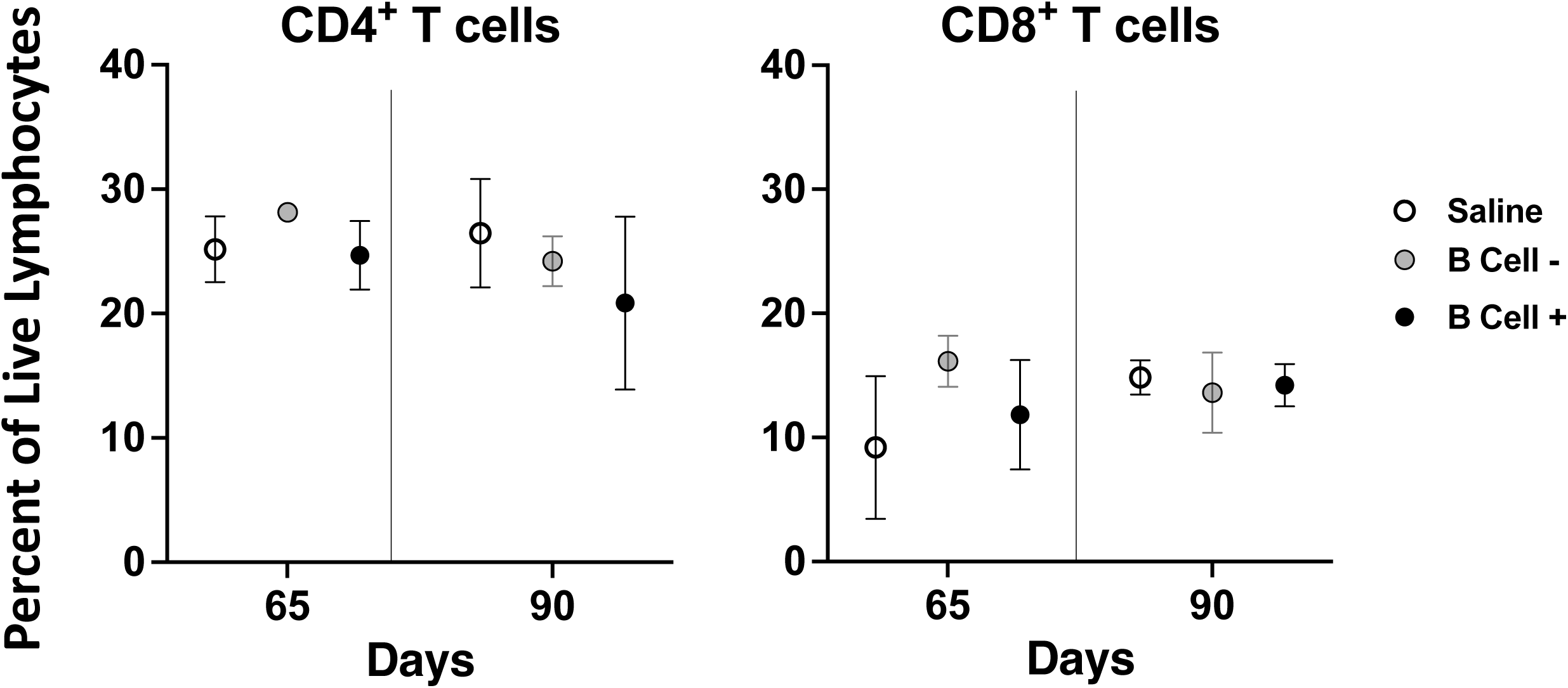
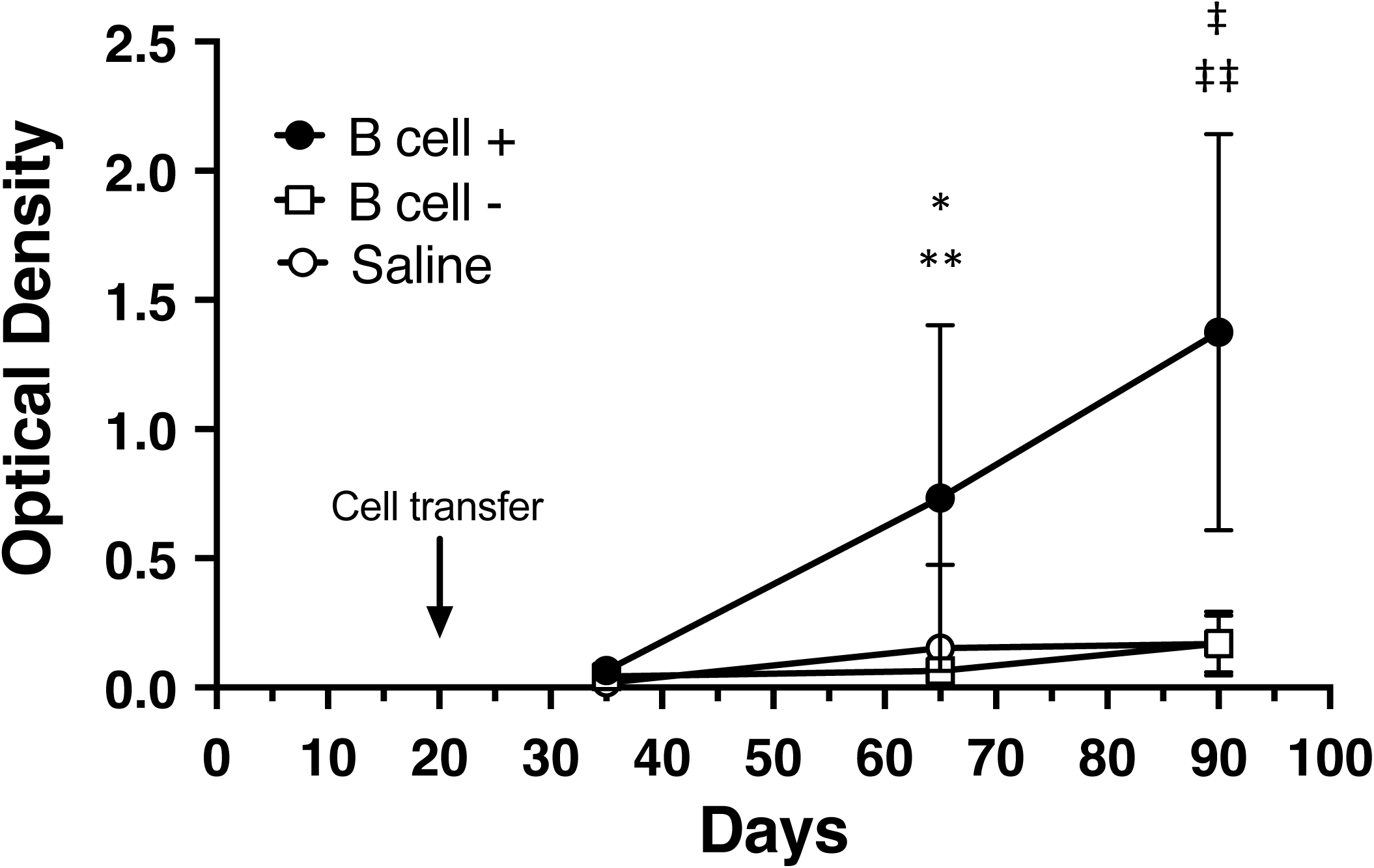
Transfer of B cells is sufficient to control *P. murina* infection in CD40 KO mice. A. CD40 KO mice were cohoused with a *P. murina* infected seeder beginning at day 0. At day 20 mice were injected via tail-vein with either PBS or ∼28 million CD19^+^ B cells from C57BL/6 mice. For A and B, organism loads were deteremined by qPCR targeting the single copy *dfhr* gene, and are expressed as *dhfr* copies per mg lung tissue. Data represent the geometric mean ± SD for 2-3 mice per time-point for the splenocyte group, and 1 (days 25 and 65) to 2 (day 90) mice per time-point for the PBS control group. Student’s unpaired t-test was used to compare the groups at each time-point. *, p=0.02 B. CD40 KO mice cohoused as above received either CD19^+^ B cells (B cell^+^, ∼50 million cells/mouse), splenocytes that had been depleted of CD19^+^ B cells (B cell-, ∼15-25 million cells/mouse) or saline at day 20. For these studies the focus was on later time-points (days 65 and 90), when the differences were seen in the prior studies, to allow for greater numbers of mice at each of these time-points, given the restrictions on the number of mice permitted per cage. In one experiment, no flow data were available due to technical difficulties at the day 65 timepoint; all animals from that time-point were included in the analysis. Data represent the geometric mean ± SD; number of mice per time-point is as follows: day 35, n=1 for each group; day 65, 7 to 12 per group; day 90, 6 to 9 per group. Student’s unpaired t-test was used to compare the different groups at each time-point. *, p=0.02 for B cell^+^ vs B cell^—^; **, p=0.02 for B cell^+^ vs saline; ‡, p=0.005 for B cell^+^ vs B cell^—^; ‡ ‡, p=0.0002 for B cell^+^ vs saline. C. CD4^+^ (left) and CD8^+^ (right) T cells, as determined by flow cytometry and shown as a percent of live lymphocytes, for the 3 groups at days 65 and 90. Symbols represent mean, error bars represent standard deviation. No significant differences were seen among the groups at either time-poiint. D. Anti-*Pneumocystis* antibodies, as determined by ELISA, for the animals in B, demonstrated that mice receiving CD19^+^ B cells developed antibodies by day 65 with a further increase by day 90, while no antibodies developed in the other 2 groups. Data represent the mean ± SD. Student’s unpaired t-test was used to compare the different groups at each time-point. *, p=0.02 for B cell^+^ vs B cell^—^; **, p=0.03 for B cell^+^ vs saline; ‡, p=0.001 for B cell^+^ vs B cell^—^; ‡ ‡, p=0.0003 for B cell^+^ vs saline.

### BMDC cell transfer provides limited control of *Pneumocystis* burden in the CD40 KO mouse model

In mice, as in humans, a major cell subset responsible for presenting antigen to T and B cells is the DC population. DCs increase their surface expression of CD40 along with other co-stimulator molecules following activation by antigen stimulation, aiding in antigen presentation and full stimulation of B or T cells. In order to elucidate the contribution of CD11c^+^ cells to control of *Pneumocystis* infection via CD40-CD40L interaction, we adoptively transferred unprimed wild-type BMDCs into CD40 KO mice approximately 3 weeks after initial exposure to *P. murina*. There was no significant difference in *P. murina* burden in the lungs of CD40 KO that received CD11c^+^ cells compared to saline alone, at any of the three time points assessed (Figure 4A). Notably, no transferred BMDCs were detected by flow cytometry in either the lungs or spleens of these mice.

**Figure 4.**
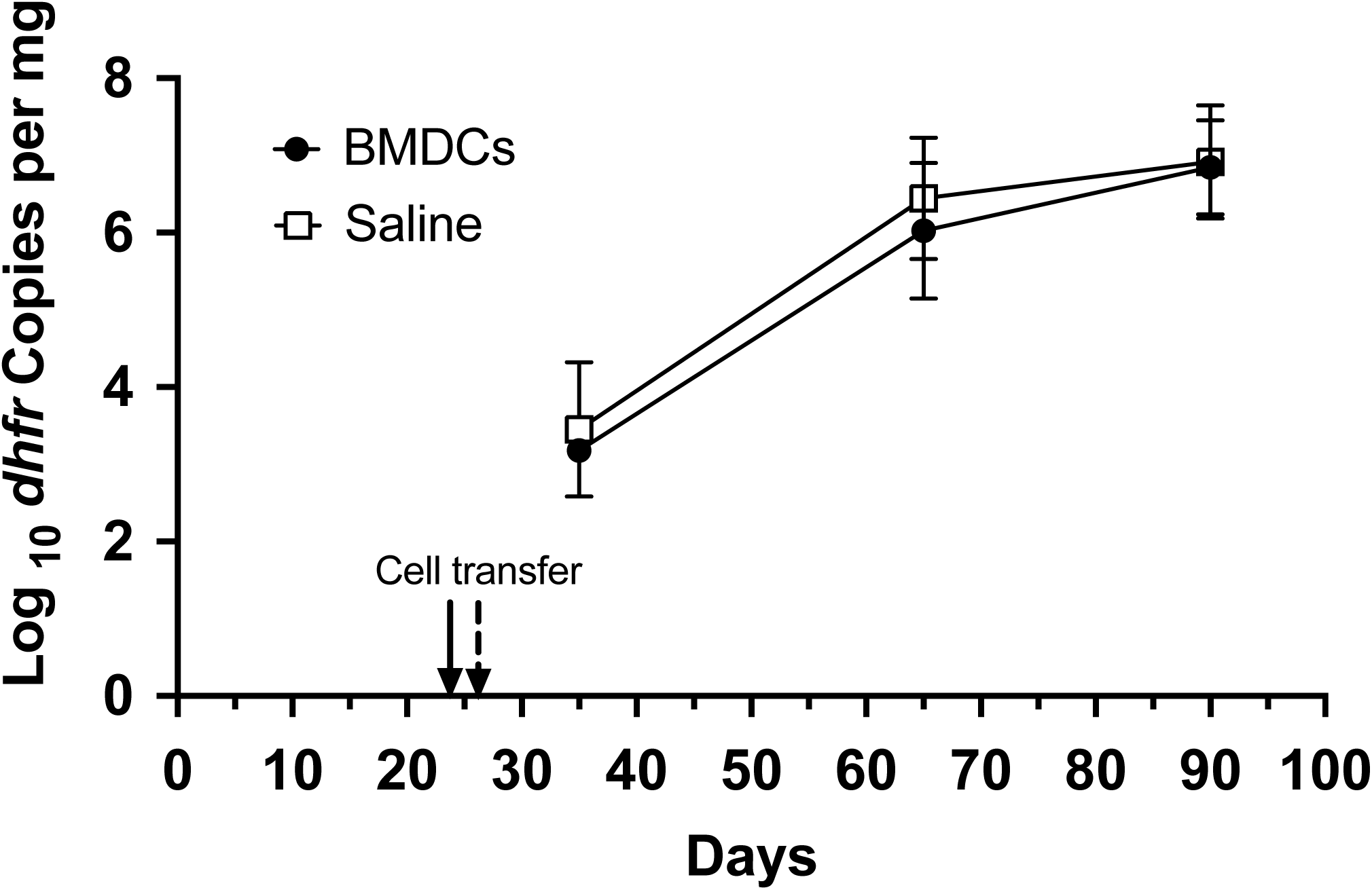
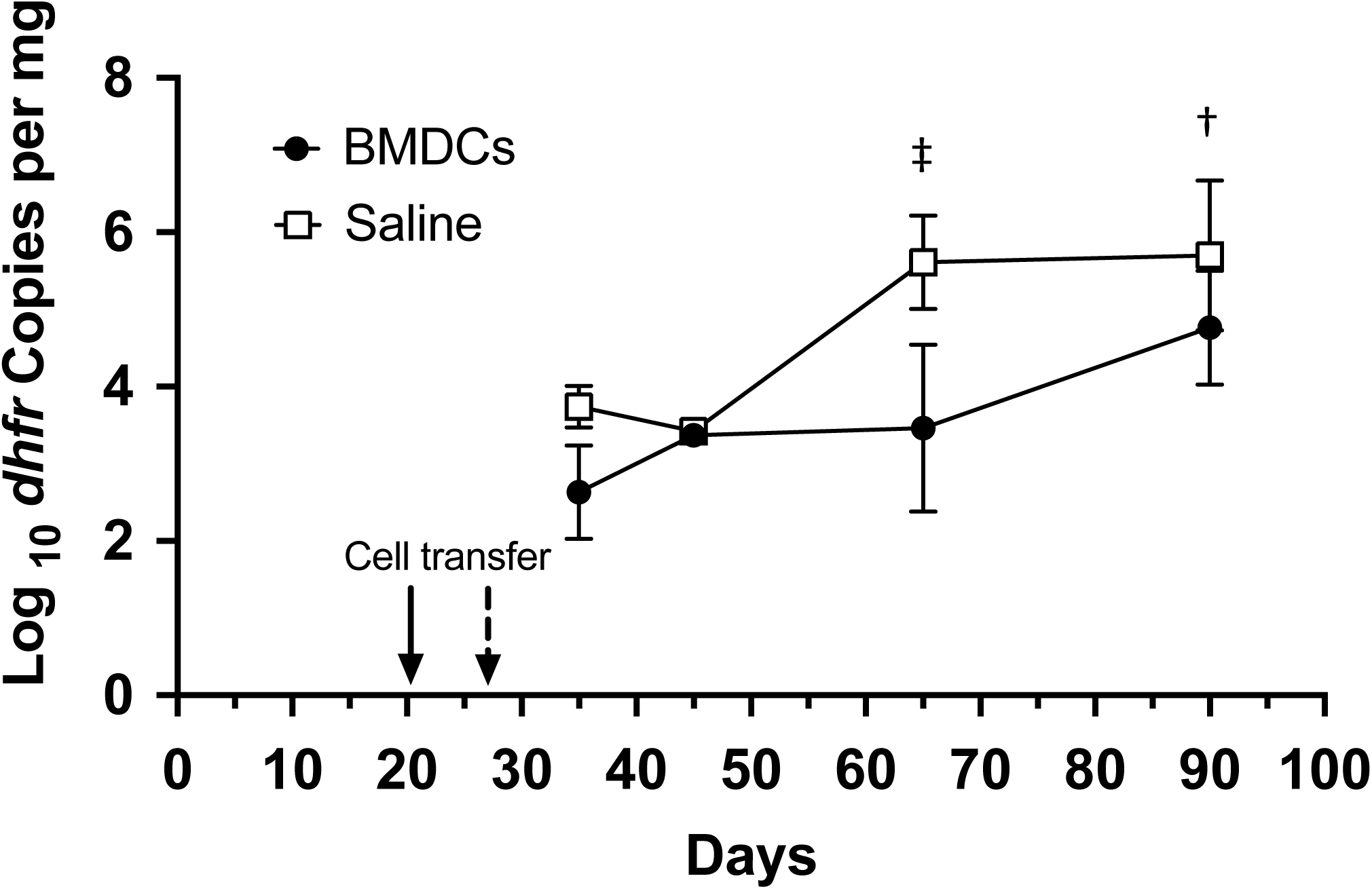
Transfer of BMDCs is insufficient to control *P. murina* infection in CD40 KO mice. A. CD40 KO mice were cohoused with a *P. murina* infected seeder beginning at day 0. At day 24-26 mice were injected via tail-vein with either PBS or ∼2.5-12 million BMDCs from C57BL/6 mice. For A and B, organism loads were deteremined by qPCR targeting the single copy *dfhr* gene, and are expressed as *dhfr* copies per mg lung tissue. Data represent the geometric mean ± SD for 2-5 mice per time-point for the BMDC group, and 2-3 mice per time-point for the PBS control group. B. CD40 KO mice cohoused as above received either PBS or 10 million antigen-primed BMDCs at day 20 (5 cages) or 27 (1 cage); the BMDCs had been incubated with a crude *P. murina* antigen overnight prior to trasfer. For the latter cage one mouse from each group was harvested at day 45 rather than day 35 as the earliest time-point. For the other times, data represent the geometric mean ± SD for 4-15 mice per time-point for the BMDC group, and 2-11 mice per time-point for the PBS control group. Student’s unpaired t-test was used to compare the groups at each time-point. †, p=0.01; ‡, p=0.001

Since antigen stimulation is critical to CD40 expression on DCs and may aid in DC trafficking, we repeated the previous experiments with BMDCs that were primed with *Pneumocystis* antigen overnight. Priming resulted in a delay in the increase of *P. murina* infection in CD40 KO mice compared to saline alone, with an ∼2 log decrease at 65 days, but only an ∼1 log decrease at day 90 (Figure 4B). Collectively, this data suggests that while CD11c^+^ DCs may contribute to the CD40-CD40L interaction associated with clearance of *Pneumocystis* infection, the contribution is modest and on its own not adequate to have a major impact on the outcome of infection in this model.

### *In vitro* antigen presentation

To determine if B cells can serve as antigen-presenting cells for *Pneumocystis* antigens, we utilized an *in vitro* proliferation assay to examine the ability of sorted CD4^+^ T cells to proliferate when incubated with a crude *Pneumocystis* antigen preparation or with purified Msg (the most abundant surface protein of *Pneumocystis* [29, 30]), either alone or when combined with purified CD19^+^ B cells, or a combination of CD11b^+^ and/or CD11c^+^ DCs and monocytes. Cells from animals immunized with adjuvant alone were unable to proliferate in response to either antigen, likely because *P. murina*-specific CD4^+^ T cells are rare in naïve mice. As shown in Figure 5, splenocytes from mice immunized with *P. murina* antigens proliferated in response to either antigen preparation. CD4^+^T cells alone were unable to proliferate to either antigen. However, there was proliferation to a crude *Pneumocystis* antigen when CD4^+^T cells were co-incubated with B cells or DCs/monocytes. Intriguingly, Msg induced proliferation when presented by DCs/monocytes, but not when presented by B cells.

**Figure 5.**
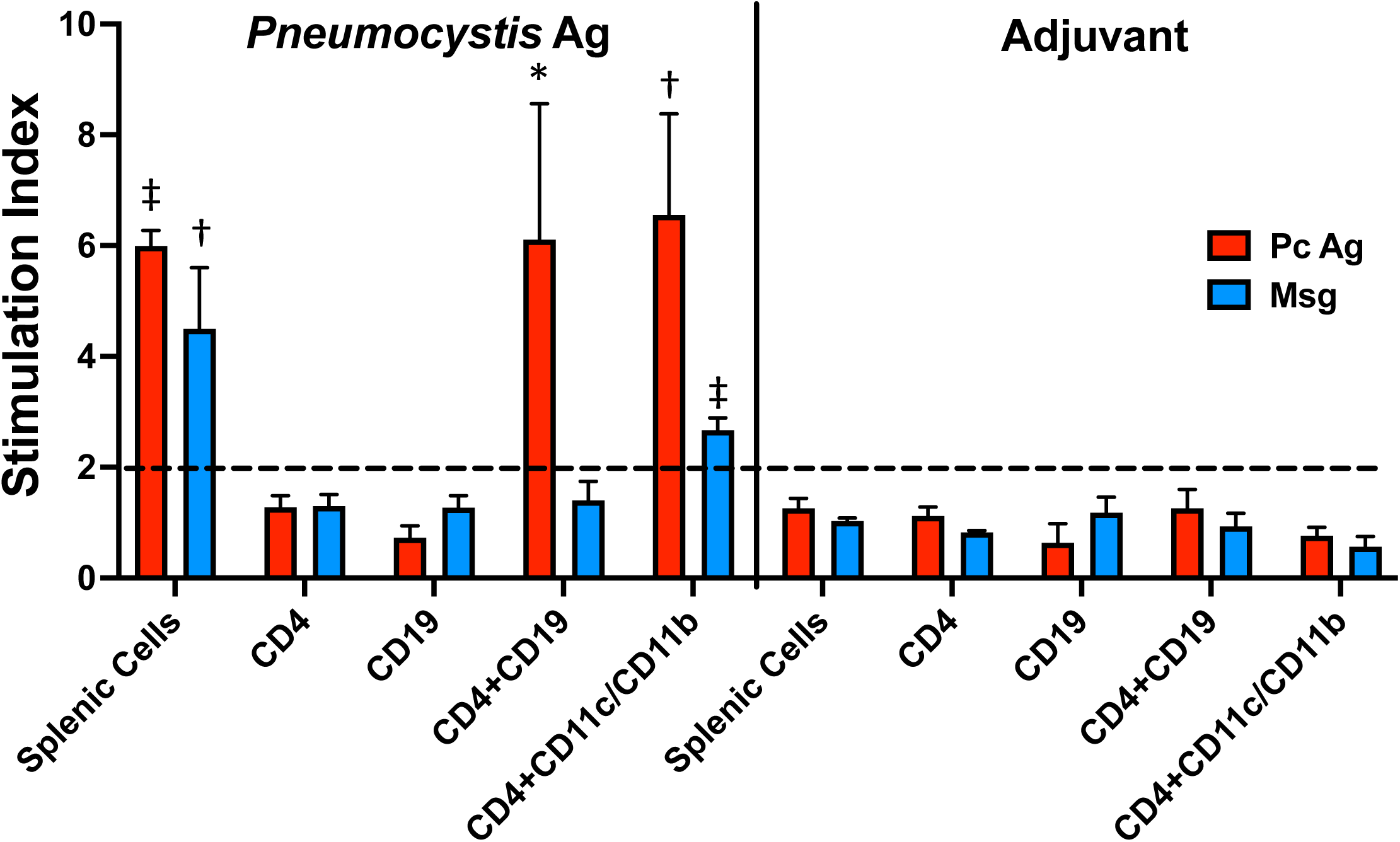
B cells can present *P. murina* antigens in vitro. A cell proliferation assay was used to determine if *Pneumocystis* antigens, presented by B cells or dendritic cells plus monocytes, could induce CD4^+^ T cells to proliferate. Spleen cells from 3-4 C57BL/6 mice that had been previously immunized with 20 µg crude *P. murina* antigen or adjuvant alone were combined and purified by cell sorting to provide populations of CD4^+^ T cells, CD19^+^ B cells, and CD11c^+^/CD11b^+^ DCs and monocytes. Purified CD4+ T cells (44,000/well) were cultured for 5 days either alone or co-incubated with B cells (50,000/well) or DCs/monocytes (5,000 cells/well) incubated with *P. murina* antigen (20 µg/ml), purified Msg (10 µg/ml) or media alone; results are shown as the stimulation index. CD4^+^ T cells or B cells cultured alone and unpurified splenocytes served as controls. Results are shown as mean stimulation index of 3 replicates for each condition, and are shown for one of 2 experiments with similar results. P values are shown for the difference between the RLU of the antigen incubated wells vs. no antigen control wells, using Student’s unpaired t-test., *p<0.05; †, p<0.01; ‡, p<0.001.

## DISCUSSION

In the current study, we have demonstrated in a mouse model that CD19^+^ B cells perform a critical role in the control and clearance of *Pneumocystis* infection, one that is mediated by CD40-CD40L interaction. The reconstitution of wild-type CD19^+^ cells in the CD40 KO mouse is sufficient to substantially control *P. murina* infection in the lungs. Notably, BMDCs did not confer the same level of control, even after priming with *Pneumocystis* antigens.

Previous studies have established that disruption of the CD40-CD40L interaction results in susceptibility to PCP both in humans and mice [3, 13, 20, 31]. We found that adoptive transfer of wild-type splenic cells was sufficient to protect CD40 KO mice from progressive *P. murina* infection. Surprisingly, however, when we utilized specifc subsets of splenocytes to clarify their roles, the decline in organism burden was associated primarily with CD19^+^ cells. The CD19^-^ splenocyte population, which would include CD40-expressing monocytes and DCs, appeared unable to control infection.

B cells have been increasingly recognized as playing an important role in controlling *Pneumocystis* infection through a mechanism that is independent of antibody production [18]. Antigen presentation to CD4^+^ T cells appears to be a major component of this mechanism, as MHC II deficient B cells are unable to reconstitute anti-*Pneumocystis* responses in a highly susceptible B cell deficient (µMT) mouse model [19, 32]. A *Pneumocystis*-specific B cell receptor appears to be required for antigen presentation, although in 1 of 2 experiments egg lysozyme specific B cells were able to substantially control infection [21]. Optimal priming of CD4^+^ T cells in the latter study required the presence of B cells during the first 2 to 3 days after infection. In addition to antigen presentation, TNF-α production by B cells may also contribute to CD4^+^ T cell activation and subsequent control of infection [22].

Given the potential importance of antigen presentation by B cells, we utilized an in vitro proliferation assay to demonstrate that B cells are able to present *P. murina* antigens to CD4^+^ T cells. This required cells from previously immunized mice, potentially because there were too few T cells recognizing *Pneumocystis* antigens in control mice. Notably, B cells induced proliferation when using a crude *Pneumocystis* antigen preparation, but not when using purified Msg, while CD11b^+^/CD11c^+^ cells were able to present both antigen preparations. The reason for these differences is uncertain, but support the concept that Msg facilitates evasion of host immune responses [27].

Anti-*Pneumocystis* antibodies play a role in limiting infection but by themselves appear insufficient to totally control/clear infection [11, 21, 33, 34]. In our studies, the decline in organism burden was associated with the production of anti-*Pneumocystis* antibodies, suggesting a role in clearance of infection; alternatively, however, this may simply reflect reconstitution of B cell function following restoration of CD40-CD40L interactions.

An important caveat for prior studies is that most utilized an intratracheal model of *Pneumocystis* infection that delivers a bolus of live and dead organisms as well as host lung cells/cell fragements that can impact the immune response. Dead and damaged *Pneumocystis* organisms will expose highly inflammatory β-glucans present in the *Pneumocystis* cysts, as well as other potential pathogen associated molecular patterns (PAMPs) which may skew responses. We have previously shown that β-glucans are largely masked in cysts in situ, which likely minimizes their inflammatory effects, and that β-glucans are major contributors to lung inflammation [35]. In the current study we have utilized a co-housing model which more closely mimics transmission and presumably immune reponses in humans [15].

Using this co-housing model, in prior microarray studies we identified a cell-mediated, interferon-γ related immune signature in immunocompetent C57BL/6 mice at approximately 5 weeks (peak of infection), followed by a robust B cell signature about a week later that persisted through 10 weeks [13]. Notably, immune reponses were essentially absent at the same time-points in CD40L KO mice despite progressive infection.

Our results using B cell depleted splenocytes as well as BMDCs suggest that other cell populations cannot replace B cells in providing optimal immunity, at least in CD40 KO mice. The limited contribution by CD11c^+^ cells to clearance of infection may have been due to ineffective trafficking of donor CD11c^+^ cells to recipient lungs. Further, BMDCs may not function identically to native DCs [36, 37]. However, based on the more prominent control seen at day 65 vs. day 90 when using primed BMDCs, we speculate that the contribution of these cells is primarily early in infection, resulting in a delay in progression kinetics, but ultimately in an inability to clear infection.

Intriguingly, in an intratracheal inoculation model, µMT mice that were reconstituted with chimeric bone marrow cells in which B cells did not express CD40, while other cells including DCs and macrophages, expressed CD40, were able to clear *Pneumocystis* infection, albeit with delayed kinetics compared to mice reconstituted with wild-type bone marrow cells [20]. This suggests that CD40 expression by B cells is not an absolute requirement, and alternative mechanisms to activate T cells are operational. Timing of reconstitution, trafficking effects, or CD40 expression by other cell populations that are reconstituted following bone marrow transplantation may account for the differences between the studies.

## Acknowledgements

This project has been funded by the Intramural Research Program of the NIH Clinical Center, National Institutes of Health. All authors report no conflicts of interest. We thank Rene Costello for providing animal care, and the NHLBI core flow cytometry facility for support of the cell sorting studies.

## Author contributions

M. S. performed and analyzed the study experiments. S.J.C., L. R. B., and Y.U. assisted with some experiments and analysis of the results. M. S. and J. A. K. conceived and designed the study. J. A. K. supervised the study. All authors contributed to the writing and revision of the manuscript.

## Notes

### Competing Interest Statement

The authors have declared no competing interest.

